# ColabFold predicts alternative protein structures from single sequences, coevolution unnecessary for AF-cluster

**DOI:** 10.1101/2023.11.21.567977

**Authors:** Lauren L. Porter, Devlina Chakravarty, Joseph W. Schafer, Ethan A. Chen

## Abstract

Though typically associated with a single folded state, globular proteins are dynamic and often assume alternative or transient structures important for their functions^1,2^. Wayment-Steele, et al. steered ColabFold^3^ to predict alternative structures of several proteins using a method they call AF-cluster^4^. They propose that AF-cluster “enables ColabFold to sample alternate states of known metamorphic proteins with high confidence” by first clustering multiple sequence alignments (MSAs) in a way that “deconvolves” coevolutionary information specific to different conformations and then using these clusters as input for ColabFold. Contrary to this Coevolution Assumption, clustered MSAs are not needed to make these predictions. Rather, these alternative structures can be predicted from single sequences and/or sequence similarity, indicating that coevolutionary information is unnecessary for predictive success and may not be used at all. These results suggest that AF-cluster’s predictive scope is likely limited to sequences with distinct-yet-homologous structures within ColabFold’s training set.

## Introduction

The authors of AF-cluster, Wayment-Steele et al., claim that ColabFold uses coevolutionary inference to predict alternative conformations of proteins from shallow sequence clusters^4^. We find no evidence supporting this Coevolution Assumption. Although coevolution of alternative states has been demonstrated for certain proteins, such as membrane proteins, it has historically been dismissed for fold-switching proteins, which remodel their secondary and tertiary structures and change their functions in response to cellular stimuli^5^. Rather, until very recently, fold-switching proteins were believed to be haphazard byproducts of evolution under no selective pressure to maintain their two distinct conformations^6^. Fold-switching proteins tend to assume a dominant easily predictable fold and an alternative fold (**Figure 1A**), whose structure is difficult to predict at least partially because of weak or absent coevolutionary signatures. These signatures were recently uncovered by aggregating signals from many multiple sequence alignments (MSAs) with hundreds to tens of thousands of sequences^7^. Curiously, Wayment-Steele et al. claim to predict alternative conformations using coevolutionary inference from MSAs with 10 sequences or fewer. Here, we analyze these MSAs and find no evidence for coevolutionary inference. Instead, we present multiple lines of evidence indicating that AF-cluster predicts fold switching from single sequences and/or by associating structures in its training set with homologous sequences, like homology modeling. This mode of operation likely limits AF-cluster’s predictive scope to sequences with distinct-yet-homologous structures within ColabFold’s training set.

**Figure 1.**
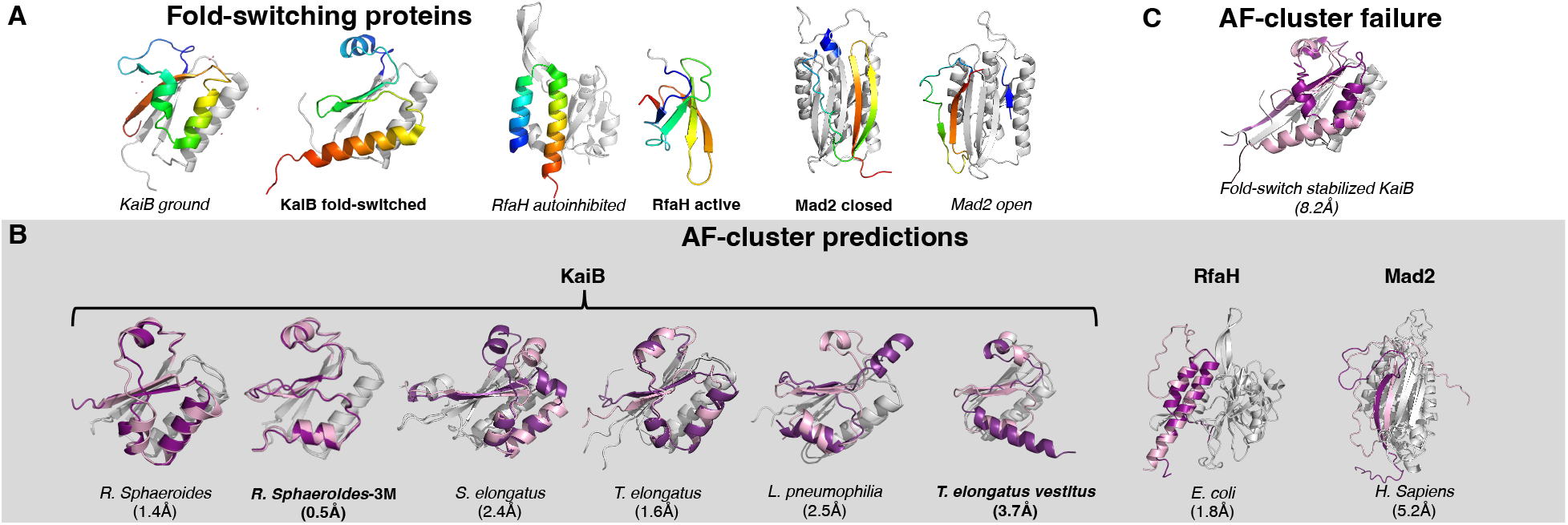
ColabFold predictions from single sequences match AF-cluster predictions. (A) Experimentally determined structures of fold-switching proteins studied by Wayment-Steele and colleagues: KaiB, RfaH, and Mad2. Fold-switching regions are rainbow, colored N-terminus (blue) to C-terminus (red); regions that do not switch folds are gray. Full-length structures of all proteins are shown, except the C-terminal domain of β-sheet RfaH is shown on its own to fit in the figure. KaiB ground often folds into a tetramer and Mad2 forms a dimer with one of each conformer. Bold labels indicate dominant (easy-to-predict) conformations, whereas italicized labels indicate alternative. (B) Superpositions of AF-cluster predictions (fold-switching regions in pink) with single-sequence predictions (fold-switching regions in purple); single-folding regions of both predictions colored light gray. All atom root-mean-square deviations (RMSDs) reported below the source organisms of each protein; bold labels indicate dominant fold prediction; others are alternative. (C) AF-cluster fails to predict the correct conformation of fold-switch stabilized KaiB, whose sequence is 92% identical to its closest homolog (T. elongatus), all-atom RMSD between most frequently predicted conformation and experiment below.

## Results

We inputted single sequences of each successfully predicted AF-cluster target into ColabFold^3^ and observed striking similarities between single-sequence and AF-cluster predictions (**Figure 1B**). All predictions were performed on sequences of experimentally characterized fold-switching proteins (**Figures 1A**). Wayment-Steele and colleagues focused most of their predictions on homologs of KaiB, whose fold switching helps to regulate cyanobacterial circadian rhythm^8^. On average, single-sequence predictions of KaiB homologs were within 2.0 Å root-mean-squared deviation (RMSD) of their respective AF-cluster predictions. The fold-switching C-terminal domain (CTD) of the transcriptional regulator RfaH^9^ predicted from a single sequence was within 1.8 Å RMSD of its corresponding AF-cluster prediction, and the C-terminal β-sheets of the mitotic spindle protein Mad2^10^, though less accurate than the other structures (5.2Å), are poised in the alternative open conformation, on the left side of the 4-β-sheet rather than the right, which corresponds to the closed dominant state. Single-sequence predictions do not result from MSA-based coevolutionary inference, casting doubt on the claim that AF-cluster leverages coevolutionary information from its shallow MSAs containing 10 sequences or fewer. These results are consistent with our previous analysis of Wayment-Steele et al.’s target Mpt53, in which we found no coevolutionary evidence for its putative alternative fold^7^. This prediction has not been confirmed experimentally. If it is found to be correct, MSA-based coevolutionary inference is likely not the explanation.

Wayment-Steele et al. mentioned that single-sequence predictions of KaiB from *T. elongatus vestitus* (KaiB TV-4) were not consistent with its experimentally observed prediction^4^. We found that was in 16/80 cases: it had a C-terminal helix rather than β-sheet. Wayment-Steele and colleagues used the secondary structure of this C-terminal region to distinguish fold switch predictions from ground. Interestingly, 6/10 of the sequences in the MSA for TV-4 generated by AF-cluster were predicted to assume the correct TV-4 conformation from single sequences. This suggested that ColabFold may use the structural inferences of these sequences to inform predictions of TV-4’s structure more robustly, analogous to homology modeling rather than coevolutionary inference. To test this possibility, we ran ColabFold with an MSA of two sequences from this cluster with correct structure predictions. Two sequences are arguably too shallow for robust coevolutionary inference. Nevertheless, we found that this MSA boosted prediction quality considerably compared to single-sequence: 4/5 predictions were within 1.8Å of the AF-Cluster prediction. This result indicates that coevolution would not necessarily explain if AF-cluster’s MSAs improve predictions, such as by requiring fewer recycling steps till the model converges. Proof of coevolution would need to rule out structural inference from sequence similarity and conservation, a difficult task with a complex neural network like ColabFold.

Furthermore, AF-cluster failed to predict the correct conformation of an experimentally characterized KaiB variant whose sequence is 92% identical to the KaiB from *T. elongatus* (**Figure 1C**). Though only 8 mutations distinguish it from its experimentally characterized ground-state homolog, this variant assumes the fold-switched state. We ran ColabFold on this sequence using the 5 AF-cluster-generated MSAs with closest average sequence identity to it. To ensure adequate sampling, we used 16 random seeds in each run, generated 5 models for each seed, and ran 3 recycles for each model. Although experimentally shown to populate the fold-switched state only, AF-cluster predicted that this variant assumes the ground state, which was predicted to within 2.5Å using all 5 MSAs. By contrast, the ground state was not predicted at this level of accuracy for any of the 5 sequence alignments. To ensure that the fold-switched state was not predicted at a lower accuracy, we searched among all 400 models and found it was predicted once at 3.5Å with low confidence (average pLDDT of 51 in fold-switching region). Together, these results demonstrate that the sequence alignments generated by AF-cluster do not always enable correct structure predictions of KaiB variants.

In their analysis, Wayment-Steele et al. used MSA Transformer^11^ to confirm the presence of coevolutionary information in some of their shallow KaiB MSAs. A broader analysis revealed lack of substantial coevolutionary information in the MSAs used in successful predictions. Here are three examples: one for the 2-sequence MSA of KaiB TV-4, which produces structures to within 1.0 Å of the 10-sequence AF-cluster prediction, one generated by AF-cluster and used to predict the structure of RfaH, and another generated by AF-cluster and used to predict the best structure of the alternative open form of Mad2. In all three cases, we see no evidence for coevolution unique to these structures (**Figure 2**), and in the case of RfaH, weak signals unique to the alternative β-sheet C-terminal domain are observed rather than the predicted α-helical conformation. MSA Transformer results are an estimate for the coevolutionary signals that ColabFold may detect. Nevertheless, Wayment-Steele et. al. use its contact maps to argue for the presence of coevolutionary information in some of their MSAs, making it reasonable to use lack of couplings to argue for their absence. This is especially true since the authors of AlphaFold2^12^– the basis of ColabFold–state that consistent prediction accuracy requires an MSA of at least 32 input sequences: more than triple the size of AF-cluster’s input MSAs.

**Figure 2.**
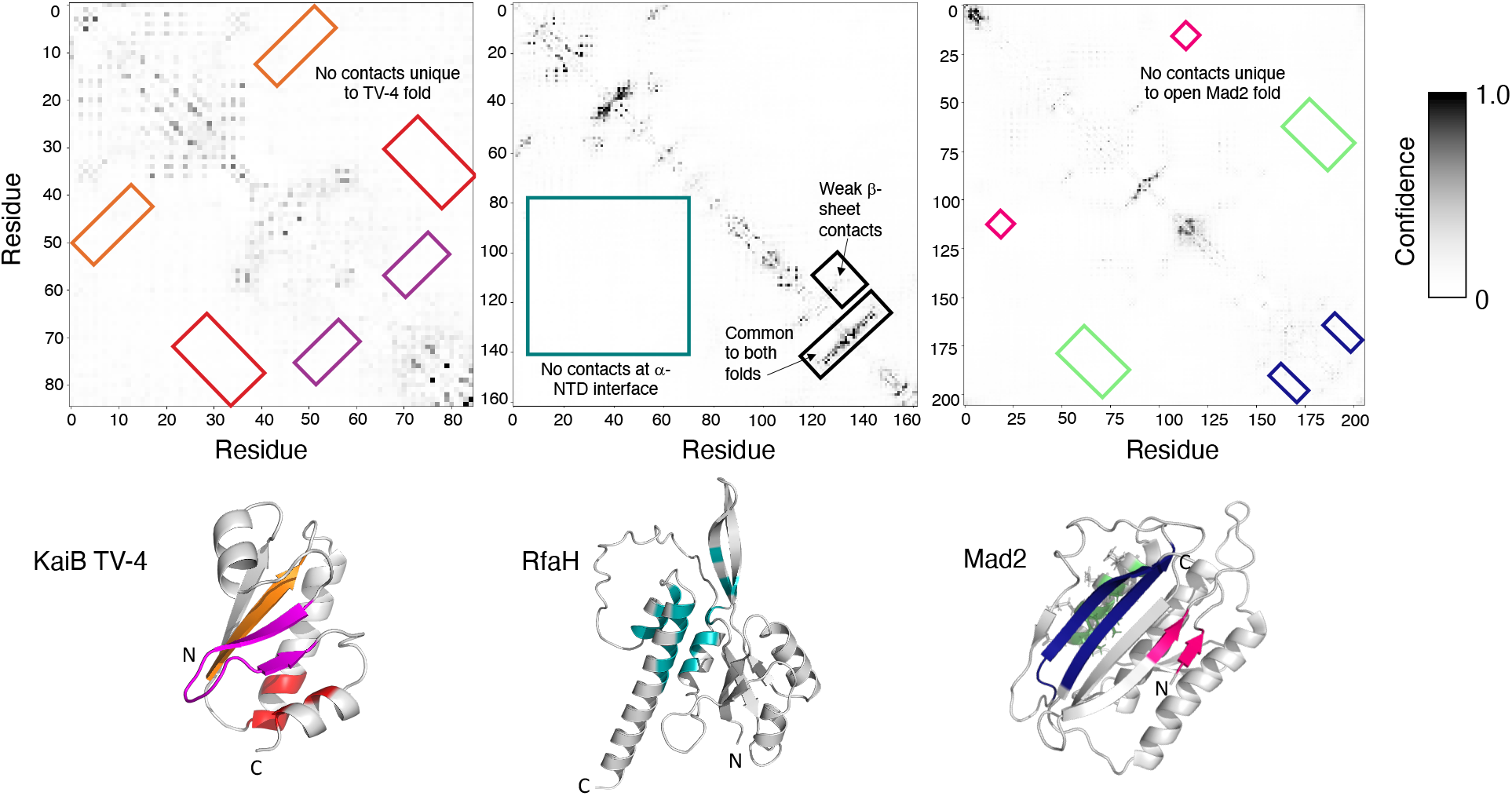
MSA Transformer predicts no unique coevolutionary signals from shallow MSAs used to predict the structures of KaiB TV-4, RfaH, or Mad2 successfully. Contacts unique to each fold are highlighted in their respective contact maps and colored according to their three-dimensional structures. Black boxes in the RfaH contact map are not represented in the structure and are used instead to annotate interesting features of its contact map.

An in-depth analysis of Mad2 provides further evidence against coevolutionary inference of alternative conformations. AF-cluster’s best prediction of Mad2’s open form originated from an MSA of four sequences with no coevolutionary evidence unique to its conformation (**Figure 2**). We used ColabFold to predict its conformation using MSAs containing each individual sequence, four sets of predictions from four MSAs, each with one sequence. In each case, 90%-100% of predictions were within 2.5Å of Mad2’s open conformation. Since successful coevolutionary inference results from combinations of sequences engendering more structural information than individuals, one would expect the MSA from AF-cluster to produce more correct predictions with greater accuracy. The opposite is true. Only 60% of predictions were within 2.5Å of Mad2’s open conformation, and their overall accuracy was substantially lower: 3.4 ± 2.2Å for AF-cluster’s MSA compared with 1.6 ± 0.2Å for the MSA with one sequence. To further investigate whether the MSA from AF-cluster might accelerate ColabFold’s convergence, we varied the number of recycle steps for ColabFold’s simulations from between 1 and 12 for both the AF-cluster and single-sequence MSAs. To enhance sampling, 16 random seeds were used for each run, and 5 models were generated from each seed. The one-sequence MSA outperformed the AF-cluster MSA by every metric after 1 recycle: it produced a greater number of accurate structures at each recycling step, converged to a higher fraction of correct predictions faster, and produced more accurate structures (**Supplementary Figure 1, Supplementary Table 1**). These results clearly demonstrate that an MSA with one sequence provides more predictive benefit for open Mad2 than the best-performing AF-cluster MSA, casting doubt on its ability to facilitate coevolutionary inference.

## Discussion

If AF-cluster is not predicting these alternative conformations from coevolutionary information, what is the basis of its predictions? As a deep-learning algorithm, the specific inner-workings of ColabFold can be difficult to explain. However, we have reason to believe that ColabFold has memorized these alternative conformations and associates them with similar sequences^13,14^, analogous to homology modeling.

AF-cluster fails to predict the alternative conformations of Selecase, Lymphotactin, and CLIC1. Nevertheless, our work has identified coevolutionary signals unique to all three of these missed conformations^7^. Interestingly, their ColabFold single-sequence predictions, unlike those of KaiB, RfaH, and Mad2, are biased toward their dominant states. One explanation is that ColabFold may not have stored information about their alternative conformations in its weights. This lack of “memorization” may better explain AF-cluster’s predictive failure than Wayment-Steele et al.’s observation that AF-cluster fails to predict oligomeric states of fold switchers, especially since it correctly predicted both the tetrameric KaiB fold and both Mad2 conformations, which dimerize.

These findings have critical implications: without coevolutionary information, ColabFold is constrained to sample alternative conformations it has “seen” during training, limiting the sorts of new fold-switching proteins it can discover. Wayment-Steele and colleagues’ results support this assertion: the alternative conformation of the novel fold switcher they propose has homologs that assume both structures that are likely in ColabFold’s training set^4^. In short, all evidence we can see points to ColabFold using sequence similarity and possibly sequence conservation patterns, but not coevolution, to make its predictions. For Wayment-Steele and colleagues to demonstrate coevolution directly, they would need use a coevolution-specific method outside of ColabFold, which combines many pattern recognition methods in its Evoformer module, making it difficult to isolate contributions from any specific MSA property (sequence similarity, conservation, coevolution, or other).

We agree with Wayment-Steele and colleagues that coevolutionary information is present in shallow MSAs with sequences similar to the target of interest. In fact, we found direct evidence of coevolution for the alternative conformations of 56/56 fold-switching proteins in subfamily-specific MSAs–including all 6 families attempted by AF-cluster^7^. These MSAs contained hundreds to thousands of sequences, not ten or fewer. In short, there is no evidence that ColabFold leverages coevolutionary information from these very shallow AF-cluster MSAs to make its predictions, which are sometimes correct for other reasons. We believe that ColabFold may use coevolution for structural inference in some of its predictions, but not alternative conformations from AF-cluster’s MSAs. The next challenge for the field is to consistently generate structural models of alternative conformations from diverse fold switchers–and eventually other proteins–by leveraging present-but-unused coevolutionary information from MSAs. The idea behind AF-cluster suggests a reasonable way forward: retrain or fine-tune ColabFold to associate different conformations with MSAs directly shown to contain corresponding coevolutionary information.

## Methods

### Single sequence predictions

Single sequences of each AF-cluster prediction were inputted into ColabFold^3^ and run in single-sequence mode with all other settings default including 5 predictions per sequence. Predictions most similar (by RMSD) to each AF-cluster prediction–taken from the AF-cluster Github repo (www.github.com/HWaymentSteele/AF_Cluster)–are shown in Figure 1. Wayment-Steele et al. mentioned that single-sequence predictions of *T. elongatus vestus* were not consistent with its experimentally observed prediction^4^. We found that they were in 16/80 cases (Colabfold single-sequence predictions with 16 random seeds, 5 models/seed, 3 recycles/model). Furthermore, running ColabFold with a two-sequence MSA (WP_011056401.1 and WP_058883586.1) improved predictions. This MSA is too shallow for coevolutionary inference, and single-sequence predictions of WP_011056401.1 and WP_058883586.1–both taken from the *T. elongatus vestus* sequence cluster on the AF-cluster Github repo–yielded the correct *T. elongatus vestus* conformation. RMSDs were calculated by comparing the full structures of proteins with their respective AF-cluster predictions, except RfaH, whose RMSD was calculated for its fold-switching C-terminal domain only. All RMSDs were calculated with PyMOL^15^.

### Contact map generation

MSA Transformer^11^ was run with default settings on a two-sequence MSA for KaiB TV-4 (WP_011056401.1 and WP_058883586.1), and the AF-cluster generated MSAs for RfaH (RFAH-049) and Mad2 (1S2H-047). Protein structures in both Figures were visualized with PyMOL^15^.

### Fold-switch stabilized KaiB

To ensure adequate sampling, we ran ColabFold with 16 random seeds in each run, generated 5 models for each seed, and ran 3 recycles for each model with the fold-switched stabilized sequence of PDB 5JYT and its 5 most similar AF-cluster MSAs as input. AF-cluster MSAs used were: WP_11056333.1, WP_00654593.1, WP_015163972.1, WP_015167870.1, WP_022607307.1. To determine sequence similarity, pairwise sequence identities were calculated between the sequence of 5JYT and all sequences within each KaiB cluster generated by AF-cluster using Biopython^16^’s PairwiseAligner. RMSDs were calculated with PyMOL^15^ referenced against the PDB structure of fold-switch stabilized KaiB (5JYT) and a model of ground-state KaiB (WP_11056333).

### Mad2 predictions

Initial Mad2 predictions were run with 2 random seeds, 5 models/seed, 3 recycles/model. When testing convergence, we ran ColabFold with 16 random seeds per run, generated 5 models for each seed with 1, 3, 6, 8, and 12 recycling steps for each of two MSAs: the best-performing AF-cluster MSA for the Mad2 open conformation (1SH2-047.a3m) and an MSA with only one sequence (F4NY50). RMSDs of each model were calculated against 3GMH, chain L using PyMOL^15^. The number of structures ≤2.5 Å were calculated and plotted in **Supplementary Figure 1**; other statistics are reported in **Supplementary Table 1**.

## Acknowledgements

L.L.P thanks Gabriel Rocklin and Brian Volkman for a helpful discussion. This work utilized resources from the NIH HPS Biowulf cluster (http://hpc.nih.gov), and it was supported by the Intramural Research Program of the National Library of Medicine, National Institutes of Health (LM202011, L.L.P.).

