## Supplementary Figure 1 for "ColabFold predicts alternative protein structures from single sequences, coevolution unnecessary for AF-cluster"

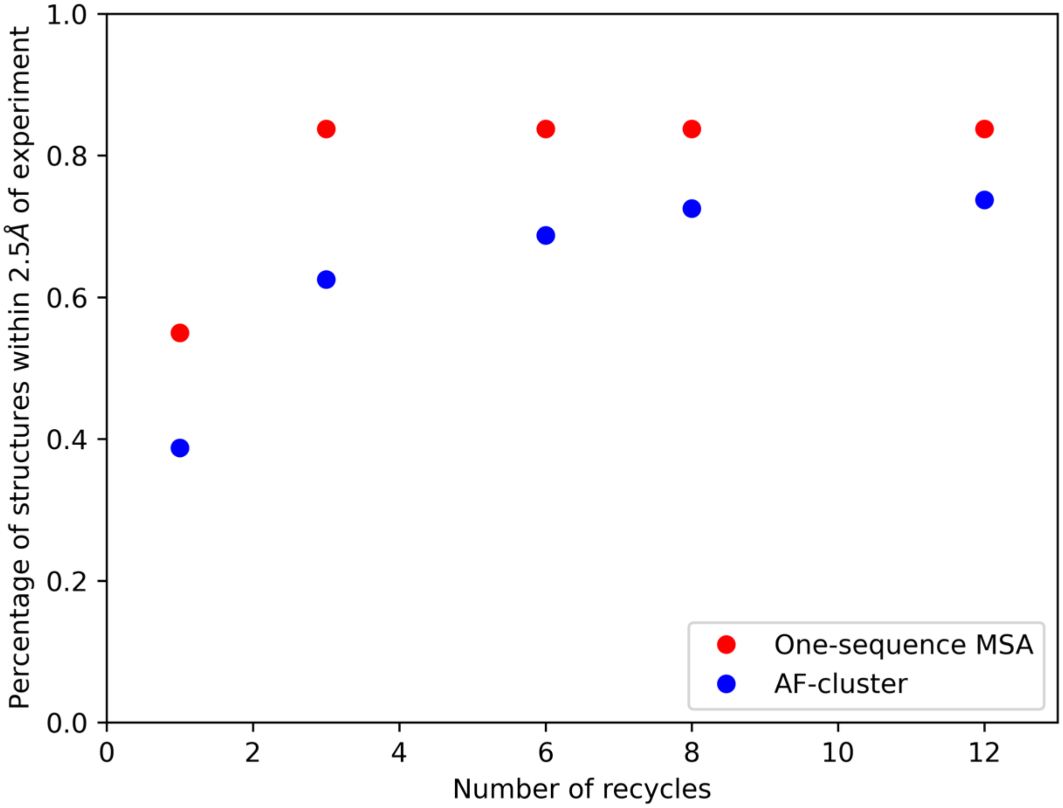


**Figure 1**. An MSA with one sequence (Uniprot ID: FNY50) enables the sequence of Mad2 to converge to the open conformation faster (with fewer recycles) than the best-performing AF-cluster MSA (1S2H-047.a3m).

| Number of recycles | Average RMSD AF-cluster | Average RMSD one-sequence MSA |
|---|---|---|
| 1 | 4.4 ± 2.6Å | 4.6 ± 4.1Å |
| 3 | 3.3 ± 2.3Å | 2.7 ± 2.7Å |
| 6 | 2.9 ± 2.1Å | 2.1 ± 2.2Å |
| 8 | 2.7 ± 2.3 Å | 2.2 ± 2.1Å |
| 12 | 2.6 ± 2.3 Å | 2.2 ± 2.1Å |

**Table 1.** An MSA with one sequence facilitates more accurate predictions of open Mad2 than the best-performing AF-cluster MSA. Errors are standard deviations from the mean. RMSDs calculated with PyMOL.
